# Inflight head stabilization associated with wingbeat cycle and sonar emissions in the Egyptian fruit bat

**DOI:** 10.1101/2020.10.15.341545

**Authors:** Jackson Rossborough, Angeles Salles, Laura Stidsholt, Peter Madsen, Cynthia F. Moss, Larry F. Hoffman

**Affiliations:** Department of Head & Neck Surgery Los Angeles, California 90095; Brain Research Institute; David Geffen School of Medicine at UCLA; Los Angeles, California 90095; Department of Psychological & Brain Sciences; Johns Hopkins University; Baltimore, Maryland 21218; Zoophysiology, Department of Biology; Aarhus University; Aarhus, Denmark

**Keywords:** gaze stabilization, acceleration, tongue clicks, motion tag, echolocation

## Abstract

Sensory processing of environmental stimuli during locomotion is critical for the successful execution of goal-directed behaviors and navigating around obstacles. The outcome of these sensorimotor processes can be challenged by head movements that perturb the sensory coordinate frames directing behaviors. In the case of visually-guided behaviors, visual gaze stabilization results from the integrated activity of the vestibuloocular reflex and motor efference copy originating within circuits driving locomotor behavior. A recent videographic study showed that echolocating bats exhibit inflight head stabilization during a target identification and landing task, though compensatory timing of the bats’ sonar signals was not reported. In the present investigation we tested hypotheses that head stabilization is more broadly implemented during epochs of exploratory flight, and is temporally associated with emitted sonar signals, which would optimize acoustic gaze. This was achieved by measuring head and body kinematics with motion sensors secured to the head and body of free-flying Egyptian fruit bats. These devices were integrated with ultrasonic microphones to record the bat’s sonar emissions and elucidate their temporal association with periods of head stabilization. Head accelerations in the Earth-vertical axis were asymmetric with respect to wing downbeat and upbeat relative to body accelerations. This indicated that inflight head and body accelerations were uncoupled, outcomes consistent with the implementation of head movements that limit vertical acceleration during wing downbeat. Furthermore, sonar emissions during stable flight occurred most often during wing downbeat and head stabilization, supporting the conclusion that head stabilization behavior optimized sonar gaze and environmental interrogation via echolocation.

**Summary statement:** Direct measurements of head and body kinematics from affixed motion sensors revealed head stabilization behaviors during exploratory flights in bats. Most sonar emissions were temporally correlated with this behavior, thereby contributing to the optimization of acoustic gaze.

## Introduction

Successful animal navigation requires sensory monitoring of selected targets and obstacles in the environment, which can be compromised by head and body movements produced by locomotion. The sensorimotor mechanisms implemented during natural behaviors typically reference head-centric coordinate frames to achieve *sensory gaze* stabilization. A prominent example of sensory gaze stabilization is the vestibulo-ocular reflex (VOR), whereby head kinematics encoded by the inner ear vestibular receptors lead to compensatory eye movements that stabilize visual targets on the retina (du Lac et al., 1995; Straka and Dieringer, 2004). Contributing to visual gaze stabilization during ambulatory activity are mechanisms to stabilize the head, thereby optimizing performance of the VOR (Dietrich et al., 2020; Dietrich and Wuehr, 2019a; Goldberg and Cullen, 2011; Shanidze et al., 2010). In birds, head movements are the predominant means of visual gaze stabilization (Land, 2015). Movements to stabilize the head can be particularly well-refined as demonstrated in a recent study showing that ultrafast head saccades were invoked for visual gaze stabilization during a rapid inflight direction reversal (i.e. “turn-on-a-dime”) maneuver executed by lovebirds (Kress et al., 2015). The head saccades were initiated most often during the start of the wing downbeat. The authors of this study interpreted this relationship as the superposition of behaviors that impair vision (i.e. visual blurring during head movements, and occlusion of the lateral visual field by the wing downbeat). Overall, this left the balance of the total wingbeat cycle with uncompromised vision (Kress et al., 2015).

While the evidence for coordinated compensatory mechanisms of eye and head movements to stabilize visual gaze have been extensively studied, comparable stabilization mechanisms for other head-centric sensory modalities have received far less attention (e.g. auditory gaze (Fischer and Pena, 2011; Genzel et al., 2016)). Of particular interest would be mechanisms of acoustic gaze stabilization in bats that utilize biological sonar to probe their environment during flight to avoid obstacles, localize conspecifics, and intercept prey (Chiu et al., 2010; Ghose and Moss, 2006; Surlykke et al., 2009). One important effector component of acoustic gaze stabilization is direction control of the sonar beam implemented by bats employing laryngeal echolocation, such as the big brown bat (*Eptesicus fuscus*; (Ghose and Moss, 2003)), as well as lingual echolocation, such as Egyptian fruit bats (*Rousettus aegyptiacus*; (Yovel et al., 2010; Yovel et al., 2011; Lee et al., 2017)). The sensory limb of acoustic gaze stabilization was recently investigated in *R. aegyptiacus* by Eitan et al. (2019), who used videographic analyses to measure head and body movements during flight. They found that head movements were strongly attenuated compared to the body center of mass during short flight segments in a target identification and landing task. This result revealed the uncoupling of head movements from oscillatory body movements associated with wingbeats, and suggested the implementation of active mechanisms for head stabilization (Eitan et al., 2019). The authors provided evidence that head stabilization was more refined as the bats approached the landing target, even while body movements were quite large. These findings suggested that head stabilization strongly depended upon mechanisms associated with echolocation-mediated target acquisition, and were not strictly driven by vestibular-mediated reflexes (i.e. comparable to oculomotor stabilization resulting from the vestibulo-ocular reflex). They also noted that the Egyptian fruit bat sonar beam elevation was directed downward upon landing, which was previously reported in a detailed study of the sonar beam pattern of this species in straight flight (Lee et al., 2017). While these data suggested a strong influence of echolocation in driving head stabilization, the temporal association of sonar emissions with head stabilization was not measured, however, which would be critical to directly establish the sensorimotor link between head stabilization and acoustic gaze. As previously described for more visually-guided species, head stabilization mechanisms likely involve multisystem compensatory reflex circuits that include vestibular reflexes and efference copy from locomotor pattern generators (Brooks and Cullen, 2019; Dietrich et al., 2020; Straka and Chagnaud, 2017; Straka et al., 2018).

The data collected by Eitan and colleagues (2019) suggest that echolocating bats may exhibit even more refined mechanisms than birds for inflight head stabilization during behavioral tasks encompassing short flights (2.5m) requiring rapid target acquisition and landing. However, the relation between head and body movements of bats engaged in exploratory flight was not investigated, representing a different behavioral state which may engage heterogeneous stabilization mechanisms. As indicated above, the timing of sonar clicks with head stabilization behavior was not recorded, precluding analyses that would have directly established correlations between these behavioral components of gaze stabilization.

In the present study we used independent multi-sensor *motion tags* that integrated triaxial accelerometers and ultrasonic microphones (Stidsholt et al., 2019) to directly record head and body kinematics and sonar clicks and test the hypothesis that Egyptian fruit bats implement head stabilization during periods of exploratory flight. We designed a similar analytical strategy as Eitan et al. (2019) focusing upon uncoupling of head and body kinematics as evidence for implementation of mechanisms to stabilize the head. The onboard ultrasonic microphone was utilized to record the bats’ inflight sonar emissions to quantify the timing of echolocation signal production in the context of head and body kinematics. In this way we also tested the hypothesis that inflight sonar probing of the environment was associated with acoustic gaze stabilization.

## Methods

### Animals

The data reported herein were collected from two Egyptian fruit bats (*R. aegyptiacus*). The female and male subjects were referred to as *Blue* and *Red,* and had body masses of 163 and 218 grams, respectively. All experimental procedures involving animals were conducted at Johns Hopkins University and conformed to the protocol approved by the JHU institutional animal care and use committee.

Inflight head and body movements, as well as the animals’ sonar emissions, were recorded as they flew across a flight room (6m X 6m X 2.5m) under infrared illumination. They were trained to locate and land on a roosting perch. Upon successful landing they were rewarded with banana and allowed to rest before being retrieved and repeating the task. Sessions lasted no longer than 30 minutes and the animals were monitored for any sign of discomfort or abnormality in their flying.

### Inflight kinematics and audio recording

Head and body kinematics during flight segments were recorded by *motion tags* (Fig. 1A) placed on the skull and interscapular region of the back, between the wings (Fig. 1B). The tags were attached with water-soluble theater glue (Hydro Mastix) to the animal’s fur and removed after each session. The motion tags, previously described in detail (Stidsholt et al., 2019), included a triaxial accelerometer, triaxial magnetometer (Kionix KX022) and an ultrasonic microphone (Knowles, FG-23329). They also included a 45mAh lithium-ion battery, on-board signal processing, and 8GB flash memory for data storage (Stidsholt et al., 2019). Each tag had a mass of 2.6 grams, and collectively (5.2 grams total load for both tags) represented 3.2% and 2.4% of the body masses for *Blue* and *Red*, respectively. These additional loads may require as much as 5% additional power to sustain flight, but would likely have minimal impact (i.e. <2.5%) on maneuverability (<2.5%; see (Aldridge and Brigham, 1988)). These devices were used in a previous study of body kinematics and direction heading in the European (common) noctule (*Nyctalus noctula*; 26 - 30 grams) and the big brown bat (*Eptesicus fuscus*; 14 – 18 grams), for which no anomalous effects of carrying the back-mounted tags on flight performance were noted for short-term deployments (Stidsholt et al., 2019).

**Figure 1.**
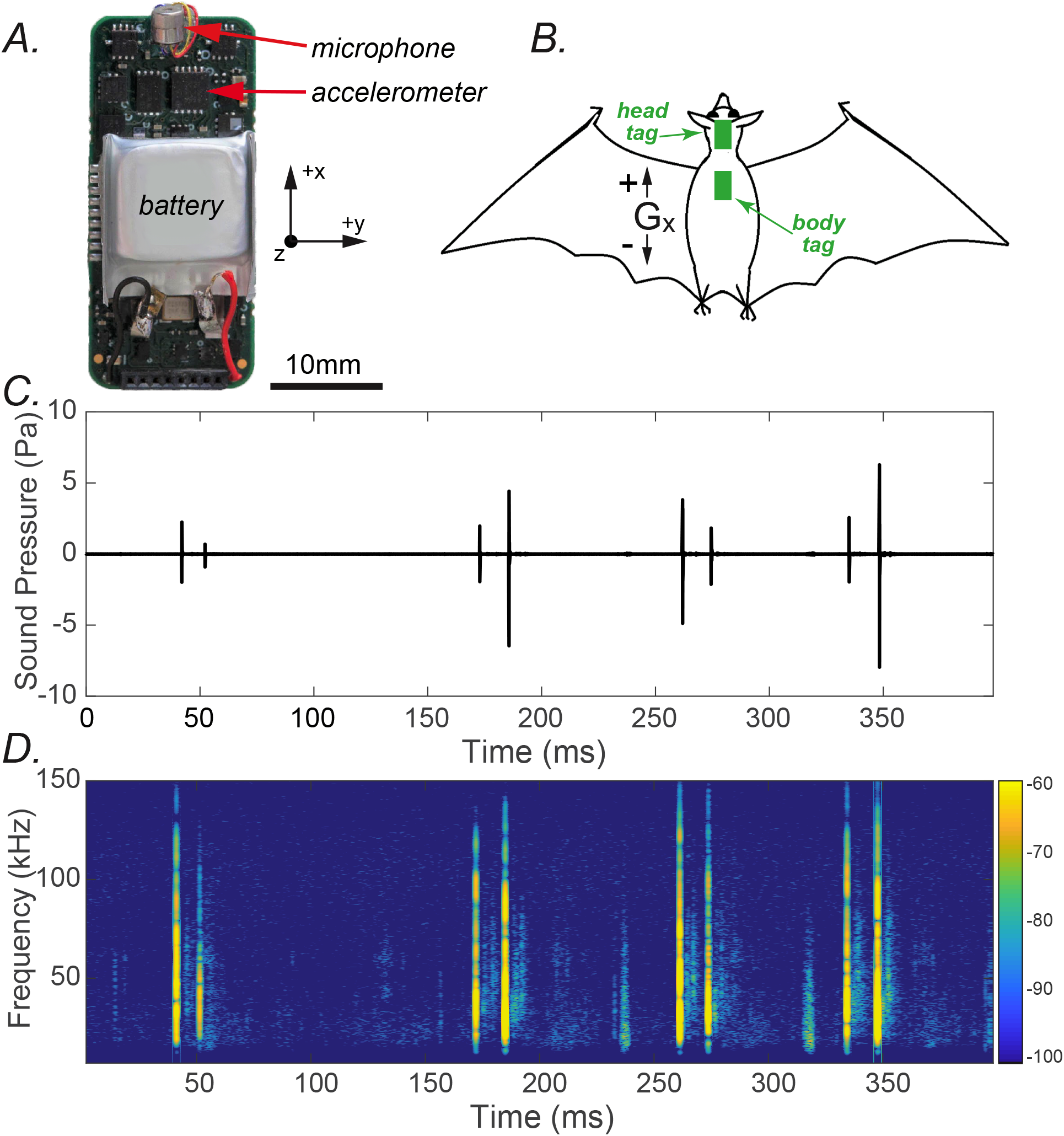
Motion tag packages placed on head and back of two R. aegyptiacus subjects for recording inflight kinematics and sonar emissions. *A*. The motion tag package was a single-board device that included a triaxial accelerometer and ultrasonic microphone. The battery is also identified. The package also included onboard signal processing and data storage (see Stidsholt et al. (2018) and text for details). The directions of the three acceleration vectors are shown by the orientation compass (+G_z_ projects out of the page toward reader). The accelerometer integrated circuit also included a triaxial magnetometer (unused in the present investigation). *B.* Diagram of *R. aegyptiacus* showing approximate placement of head and body tags. The direction of Gx is shown to confirm orientation of the tags relative to body axes. *C.* The filtered output of the ultrasonic microphone clearly shows the doublet pattern of tongue clicks emitted by *R. aegyptiacus*. The spectrogram of this emitted recorded segment is shown in *D*, illustrating that the doublets are composed of audio frequencies exceeding 100kHz. Spectrogram intensity scale in units of dB/kHz.

The triaxial accelerometer was oriented on each animal to align the tags’ G_x_ axis and board with the long axes of the animals’ bodies (Fig. 1A and B; see compass). Its data acquisition specifications featured 16-bit sampling at 1k samples/s for each channel, and onboard processing included a 250Hz 4-pole anti-aliasing filter. These data were downsampled offline to produce accelerometer measures at 100 samples/s.

Sonar tongue clicks emitted during flight were recorded by the ultrasonic microphone sampled at 187.5kHz (Fig. 1A, 1C, and 1D). Onboard processing of microphone recordings included a first stage 80kHz 4-pole anti-aliasing filter, followed by a second filtering stage (i.e. 10kHz, 1-pole high-pass filter) to reduce wind and wing noise. This processing strategy was sufficient for recording the broadband clicks emitted by *R. aegyptiacus* that typically exhibit peak energy at approximately 35kHz (Lee et al., 2017). This preprocessed audio channel was then digitized at 16-bits and stored in flash memory. Further offline processing (described below) was implemented to unambiguously identify the ultrasonic tongue clicks emitted by the *R. aegyptiacus* subjects (Fig. 1C-D).

### Analyses of inflight head and body kinematics

#### Synchronizing head and body tags

In order to compare the acceleration data collected by head and body motion tags it was necessary to temporally synchronize the recordings. This was achieved by presenting an external audio signal that could be recovered in the respective audio channels of each tag. The audio records from both motion tags were aligned on this recorded signal. The onboard microcontroller software implemented a 50msec delay in recording data from the accelerometer, necessitating a corresponding time shift in the accelerometer data channels, but were otherwise temporally synchronized and sampled by the same onboard clock.

#### Selection and analysis of stable flight periods

An example of inflight body kinematic behavior of one *R. aegyptiacus* subject (*Red*) is shown in Fig. 2, illustrating for one subject the episodic periods of flight that was exhibited by both animals in the flight room. Among all flight episodes, those selected for analysis exhibited a minimum duration of 1.6 seconds. The first and last 2-3 wingbeat cycles of each flight were omitted to avoid take-offs and landings, thereby identifying periods of “stable” flight in the middle of each episode. The *Matlab* function *findpeaks()* was used to identify the peaks and troughs of head and body G_z_ for each flight epoch that conformed to specific criteria for minimum inter-peak/trough period (7 samples, or 0.07sec) and minimum magnitude (0.5g). Troughs were identified by inverting the acceleration epoch polarity and applying the *findpeaks()* function, in these cases utilizing a minimum magnitude of 0.2g. A total of 25 flight epochs were analyzed from each subject, comprised of 417 and 340 wingbeat cycles for *Blue* and *Red* subjects, respectively. These flight epochs were utilized for all analyses, including those of the temporal correlation of tongue click onset and G_z_.

**Figure 2.**
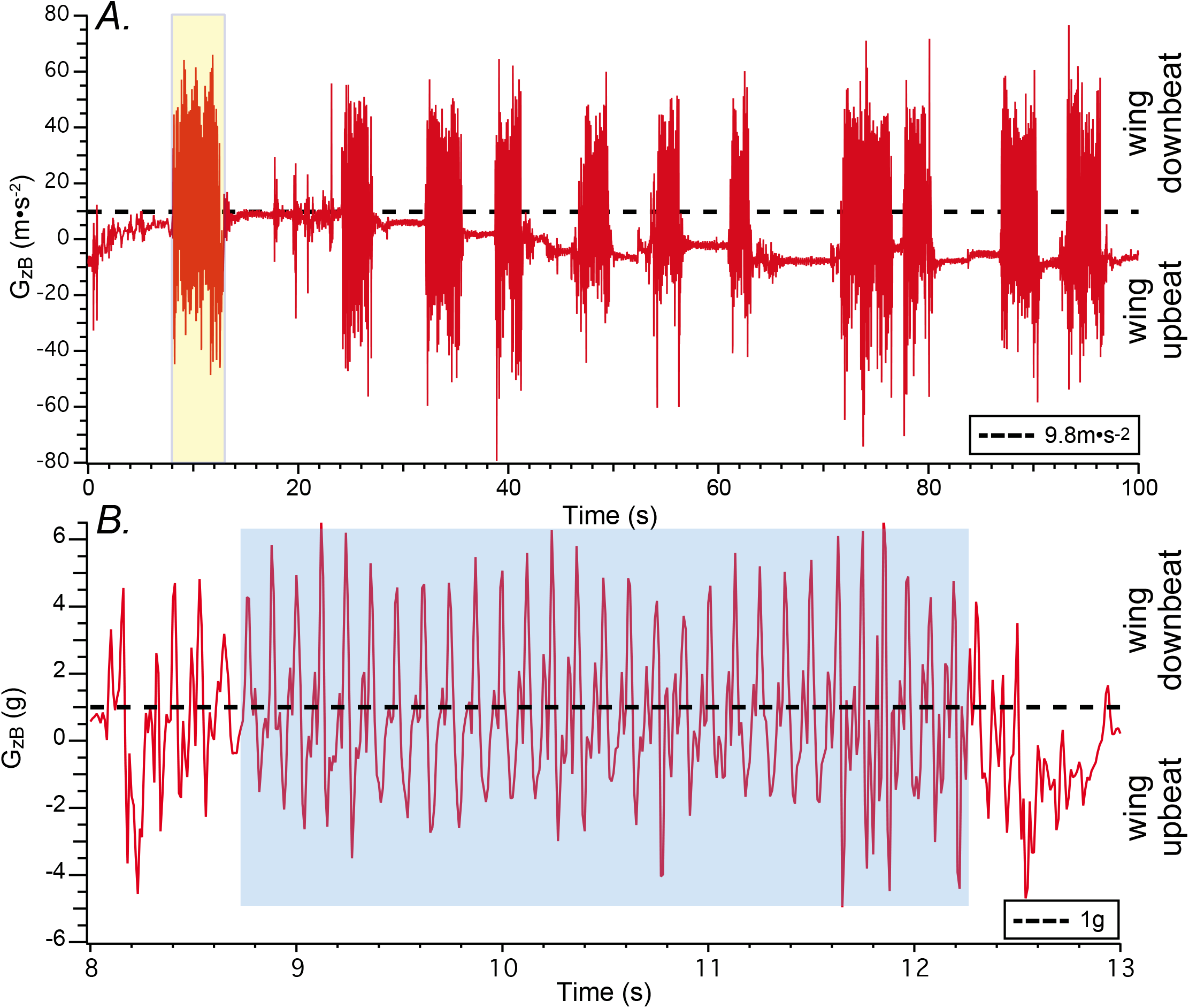
Laboratory flights comprised of short epochs. *A*. Episodic flights of one *R. aegyptiacus* subject (*Red*) within the JHU flight room represented by modulation of acceleration in the *z*-axis of the body motion tag. Wing downbeats result in an upward (antigravity) force that is greater than Earth gravity (9.8m•s^−2^; dashed horizontal line), while wing upbeats generate force vectors in the opposite direction. The first epoch (yellow highlight) is shown at higher temporal resolution in *B*. *B.* Expanded temporal view of G_zB_ highlighted flight epoch from *A*. Within this epoch a subsegment was selected for detailed analysis representing stable flight, avoiding short periods after takeoff and before landing. Acceleration magnitudes were converted to *g* (9.8m•s^−2^/g). Maximum downbeat acceleration magnitudes (*peaks*) range from approximately 4 – 6g, while maximum upbeat acceleration magnitudes (*troughs*) range −2 - −4g.

### Accelerometer data

All accelerometer data used in the present analyses were calibrated by aligning the z axes of each motion tag parallel with Earth gravity, representing 9.8m•s^−2^, and then converting to *g* (1*g* = 9.8m•s^−2^) to produce axial acceleration of G_z_ for head and body tags (i.e. G_zH_ and G_zB_). Acceleration data reflect the direct measures from each tag.

### Analyses of inflight audio

#### Identification of tongue clicks

Audio recordings during the flight epochs selected for head and body kinematic analyses were analyzed for sonar emissions. A 4-pole high-pass Butterworth filter (15kHz corner frequency) was used to remove low frequency components not associated with *R. aegyptiacus* tongue clicks (Griffin et al., 1958). It was implemented in *Matlab* using the *filtfilt()* function, a digital filter function that did not induce phase shifts or temporal delays in the resulting audio signal. Echolocation clicks emitted by the bats could then be readily identified as high frequency transients in the recorded audio channel (Fig. 1C and D). The *Matlab* function *findpeaks()* was used to locate these transients in the audio recordings with interpeak times of that ranged 16 - 28 msec and exceeded a minimum amplitude threshold of 0.75 Pa.

#### Head kinematics at click onset

Previous studies have suggested that inflight head stabilization behaviors in birds are executed to optimize visually-guided flight. A comparable behavioral paradigm in echolocating bats would be supported by the association of head stabilization with the onset of sonar emissions to optimize acoustic gaze. To evaluate this association, we determined G_zH_ and G_zB_ magnitude at the onset of sonar emissions, represented by the first emission of the tongue click doublet. Since the sampling frequency of the acoustic recording was orders of magnitude greater than that of the acceleration channels, G_zH_ and G_zB_ magnitudes at first click times were determined by linear interpolation. This was achieved by estimating the acceleration magnitude at the precise click onset time using the *Matlab* function *interp1()* from measured acceleration values immediately before and after these emission times.

### Statistical analyses

Similar to the strategy utilized by Eitan et al. (2019) in their evaluation of the angles between flight direction (determined from body position) and target, and between head position and target, head stabilization behaviors were interpreted to be represented by the uncoupling of head and body accelerations along the z-axis (i.e. G_z_) during portions of each wingbeat cycle. The present dataset utilized the distributions of head and body acceleration measures provided by the motion tags, from which temporally associated maxima in positive and negative values (i.e. acceleration peaks and troughs) comprised new distributions. In most cases, these distributions were compared by computing the Kullback-Leibler divergence (D_KL_; (Mackay, 2003)), representing measures of relative entropy. The general strategy was executed through evaluations of null hypotheses that the distributions subject to comparison were similar, which were achieved by determining the probability that the empirical D_KL_ (i.e. computed from the empirical distributions) could have resulted from random sampling from all pertinent acceleration measures. This involved bootstrap resampling from the respective distributions under evaluation. That is, D_KL_ values were computed from randomly resampled distributions, and repeated one million times to generate a null distribution of D_KL_ measures. The probabilities that the empirical D_KL_ value could have been achieved from random resampling of acceleration measures were then determined and reported.

## Results

### Body G_z_ and identification of “stable” flight

The brief, episodic flights typical for bats in a laboratory flight room environment are shown in Fig. 2A (e.g. (Stidsholt et al., 2019)). These data represent the calibrated recording of G_z_ from the body motion tag (G_zB_ accelerometer axis orthogonal to the tag’s long axis; Fig. 1A). For the *R. aegyptiacus* subjects of the present study individual epochs ranged 2 – 4 seconds in duration. These oscillations are the result of wing downbeats generating accelerations against Earth gravity (>9.8m•s^−2^), and wing upbeats generating accelerations coincident with Earth gravity (<9.8m•s^−2^). Hence, the oscillations in G_zB_ were a direct reflection of wingbeats, offset by variation in the orientation of the body tag relative to Earth gravity (e.g. when the animal lands). That is, when the *z* accelerometer axis is normal to Earth gravity G_zB_ = 0 (e.g. Fig. 2, time = 36 – 38 seconds).

Fig. 2B shows the highlighted flight segment from Fig. 2A (yellow) after expanding the time axis to illustrate G_zB_ accelerations with each wingbeat. Most wingbeats show greater peak G_zB_ acceleration during wing downbeat than upbeat, consistent with previous descriptions of the heterogeneous forces corresponding with each wingbeat half-cycle (Aldridge, 1987; Hedenstrom and Johansson, 2015). The shaded region (blue) exemplifies the *stable* portion of the flight epoch selected for analysis of head and body kinematics that excluded the onset and termination portions exhibiting more unstable accelerations associated with takeoff and landing. The analyzed flight epoch shown in Fig. 2B was approximately 3.4 seconds and included 28 wingbeat cycles. The mean period for these wingbeats was 0.121 sec, corresponding to a mean frequency of 8.3 Hz. In fact, the median wingbeat period for each subject over all analyzed flight epochs was 0.12 seconds. This is comparable to previously reported measurements of wingbeat period and frequency for *R. aegyptiacus* (Yartsev and Ulanovsky, 2013).

### Heterogeneity in the peaks and troughs of G_zH_ and G_zB_ suggests kinematic uncoupling

The record of G_zB_ in Fig. 2A illustrates that measures of acceleration reflect both dynamic (i.e. due to wingbeats) and static (i.e. due to changes in orientation affecting Earth gravity sensing) forces to produce composite measurements of head and body acceleration along the G_z_ axis. We evaluated G_zH_ in the context of G_zB_ for evidence that the head was either coupled or uncoupled from the body. Evidence that the head and body were consistently uncoupled over the 757 wingbeat cycles in both subjects would strongly infer the implementation of specific behaviors to stabilize the head during flight.

The data in Figs. 3A and B demonstrate a consistent asymmetry in the peaks and troughs of G_zH_ relative to G_zB_ highlighted by the symbols marking these local (i.e. wingbeat cycle-by-cycle) positive and negative maxima in G_z_. That is, the differences between peak G_zH_ and G_zB_ appear to be greater than the differences between the troughs of G_zH_ and G_zB_. These differences are illustrated by box-and-whisker plots in Fig. 3C and D, representing the distributions of all G_zH_ and G_zB_ measures at the acceleration peaks and troughs for *Blue* and *Red* subjects (respectively). These analyses confirm that the relationships demonstrated in the 2-second flight raw data (Fig. 3A and B) were consistently observed over the 757 wingbeat cycles in the analyzed flights of both subjects. That is, minimum G_zB_ is modestly lower than minimum G_zH_ where median G_zB_ is approximately equal to the 25^th^ quartile of G_zH_. However, G_zH_ exhibits a greater difference from G_zB_ at the peak acceleration (peak G_zH_ and G_zB_ during wing downbeat). In view of the similarities across each subject these data were combined in Fig.3E showing box-and-whisker plots of the differences between G_zH_ and G_zB_ at the maxima and minima. These values were computed as the differences in absolute values between G_zH_ and G_zB_ for both trough and peak accelerations in each wingbeat cycle. The data show that the differences in G_zH_ and G_zB_ for peak maxima appear to be larger than the differences in trough G_zH_ and G_zB_.

**Figure 3.**
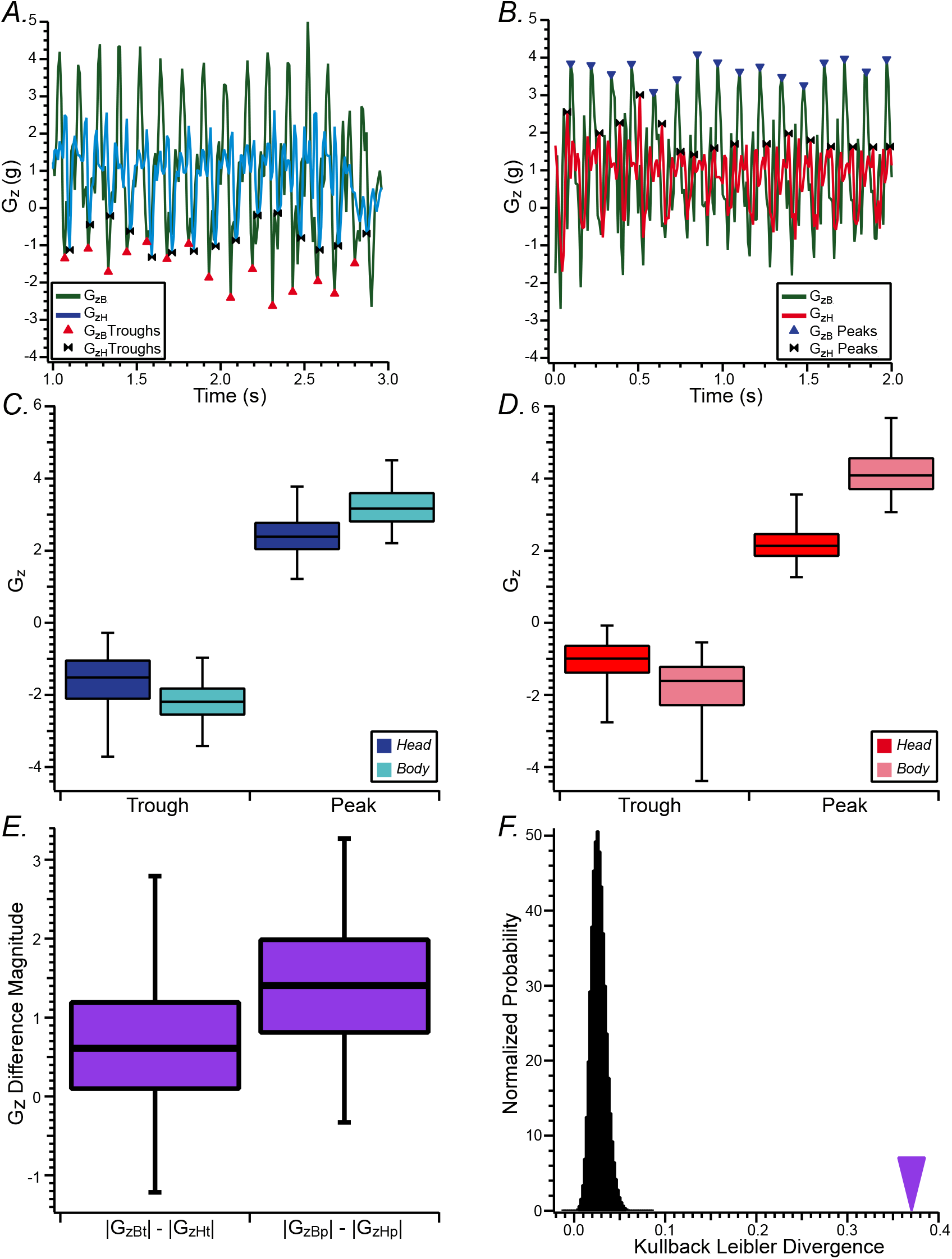
G_zH_ maxima associated with wing downbeat (peaks) and upbeat (troughs) were attenuated relative to corresponding G_zB_ maxima, and exhibited peak and trough asymmetries. *A, B*: Representative G_zH_ and G_zB_ for *Blue* (A) and *Red* (B) *R. aegyptiacus* subjects for two-second flight epochs, illustrating the identification of troughs (see marker legends in *A*) and peaks (see marker legends in *B*). These records illustrate that the differences in G_zH_ and G_zB_ peaks are greater than the differences in G_zH_ and G_zB_ troughs. *C, D:* The distributions of G_zH_ and G_zB_ peaks and troughs for *Blue* (*C*) and *Red* (*D*) subjects are represented as box-and-whisker plots, further illustrating that the differences between G_zH_ and G_zB_ peaks are greater than the differences between G_zH_ and G_zB_ troughs. *E.* For each cycle, the differences in absolute values between trough and peak G_zB_ and G_zH_ measure were obtained, and the distribution of these differences are represented in box-and-whisker plots. The D_KL_ was determined for the empirical distributions, and was compared to D_KL_ values computed from 10^6^ resampled distributions to test the null hypothesis that the empirical value of D_KL_ could be derived from random resampling of G_zB_ and G_zH_ trough and peak differences. *F.* The empirical D_KL_ value is quite different from the 10^6^ D_KL_ values obtained from random resampling, indicating that the probability of obtaining the empirical D_KL_ from random resampling the trough and peak differences is less than 10^−6^. This result supported rejection of the null hypothesis.

We compared the empirical distributions in peak and trough differences between G_zH_ and G_zB_ (Fig. 3E) by computing D_KL_, then tested the null hypothesis that this empirical value could have resulted from D_KL_ values computed from randomly resampled distributions of peak and trough differences in G_zH_ and G_zB_. The results of this analysis are shown in Fig. 3F, where the distribution of D_KL_ computed from 10^6^ randomly resampled distributions (i.e. virtual differences in peak and trough G_zH_ and G_zB_) is shown as the histogram at left and the D_KL_ value computed from the empirical distribution of peak and trough differences is shown at the arrowhead. This analysis demonstrates that the probability of the empirical D_KL_ value arising from randomly sampled difference measures is less than 10^−6^, strongly refuting the null hypothesis. This supports the conclusion that inflight peak G_zH_ exhibited a greater difference from peak G_zB_ than the differences in trough G_zH_ and G_zB_, representing an asymmetry for each wingbeat cycle.

The framework for interpreting relative head and body acceleration magnitudes along G_zH_ and G_zB_ axes is illustrated in Fig. 4, providing simple models of representative conditions in which head and body are coupled (Figs. 4A-C) or uncoupled (Fig. 4D, E). The relative head and body positions are depicted by the “stick” figures, and the motion tags are represented by the red and green rectangles affixed to head and body, respectively. In each case shown in Fig. 4 head and body accelerations result from relationships depicted by the stick figures throughout each wingbeat cycle. Wingbeat-driven modulations of G_zH_ (red) and G_zB_ (green) are modeled as 8Hz sinewaves (0.125s period) in the plots of G_z_ vs. time associated with each stick figure. G_zH_ and G_zB_ amplitudes are biased by their orientations relative to Earth gravity and the anticipated changes through the wingbeat cycle.

**Figure 4.**
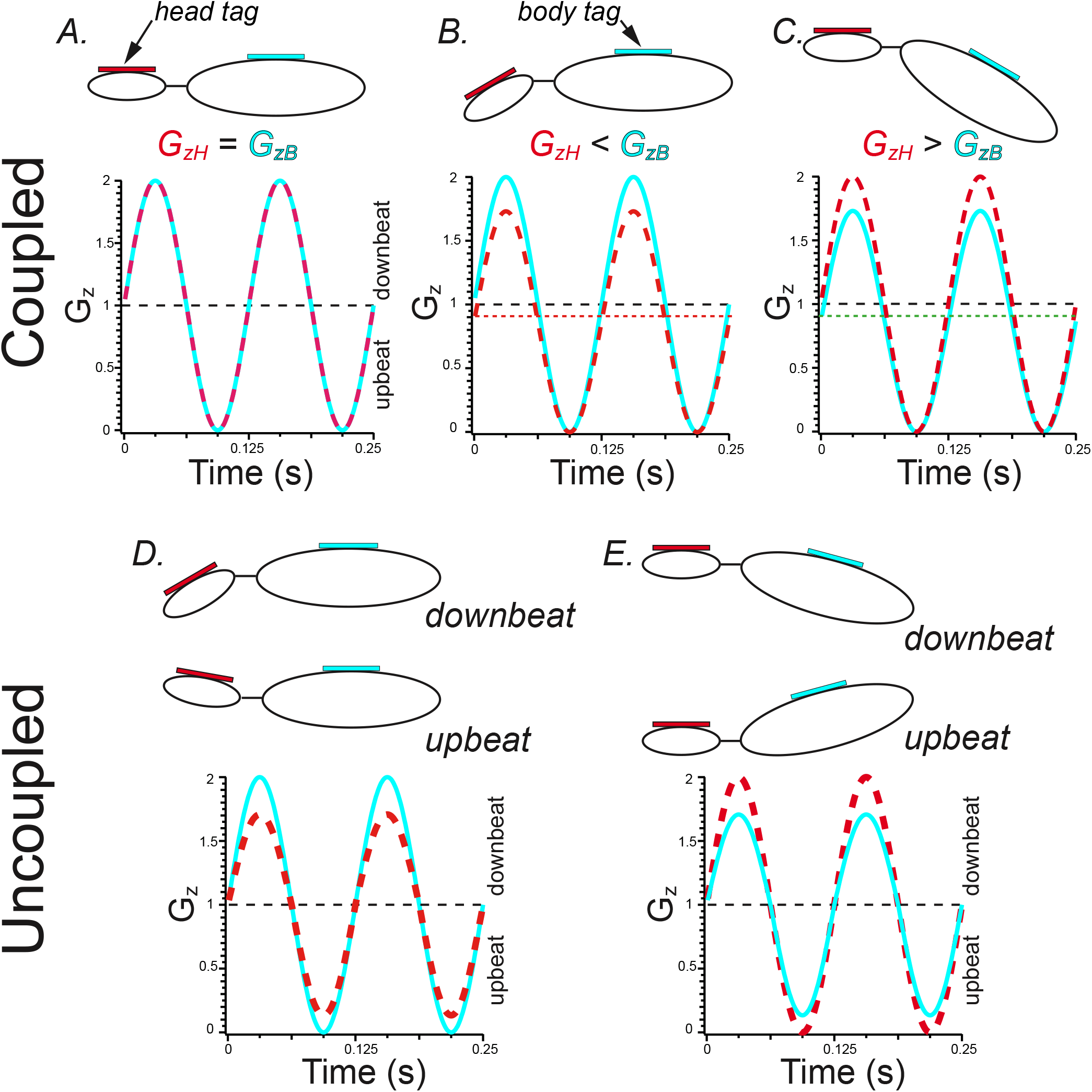
Model of G_zH_ uncoupling from G_zB_ consistent with empirical asymmetry in G_zH_ peak and trough asymmetry. *A-C*: The coupling of head and body kinematics results in fixed relationships between G_zH_ and G_zB_ measures that depend upon the orientation of the motion tags relative to Earth-gravity and the wingbeat-associated acceleration vector. Note the fixed offsets (attenuated static gravity measure) for G_zH_ and G_zB_ for conditions *B* and *C*. *D, E.* Uncoupling of head and body results in asymmetric differences in peak and trough G_zH_ and G_zB_. The configuration represented in *D*, resulting from changes in head orientation, are consistent with the empirical asymmetries shown in Fig. 3.

Three conditions in which head and body kinematics are coupled are shown in Fig. 4A-C. The characteristic features of these conditions are the fixed relationships of acceleration magnitudes at peak and trough G_zH_ and G_zB_. Fig. 4A represents the condition in which head and body motion tags are orthogonal to Earth gravity and wingbeat modulation of G_z_, resulting in comparable amplitudes of G_zH_ and G_zB_ at both maximum wing downbeat (peak) and wing upbeat (trough). The condition represented in Fig. 4B is one in which the body tag is orthogonal (i.e. optimally oriented) to Earth gravity and wingbeat-generated acceleration along the G_z_ axis, but the head is pitched down relative to the body. In this orientation the head tag is not optimally oriented for Earth-vertical sensing and therefore the acceleration magnitudes are less than that registered by the body tags. Note that wingbeat-generated G_zH_ is modulated about a slightly reduced static magnitude due to head pitch. Since head and body are coupled, the magnitudes in maximum downbeat and upbeat G_zH_ are similarly attenuated compared to G_zB_, which is positioned orthogonal to Earth gravity in this model. Due to the attenuation in Earth-gravity sensing of the head tag (i.e. due to static head pitch), peak upbeat (trough) G_zH_ and G_zB_ are similar. This attenuation in G_zH_ is not exclusive to nose-down head pitch, and would be similar in a nose-up head pitch. The third example of head and body coupling (Fig. 4C) depicts the condition in which the head is orthogonal and optimally positioned to detect Earth gravity and the wingbeat acceleration vectors in G_z_, but the body is pitched (either up or down). In this condition the modulation in G_zH_ exceeds G_zB_, but due to the reduction in static gravity sensing (dashed green line) maximum upbeat acceleration for both head and body are similar.

The conditions in which head and body are uncoupled during flight are shown in Figs. 4D and 4E, illustrating the changes in head position relative to the body during down- and upbeat portions of the wingbeat cycle that are consistent with head and body uncoupling during flight. In the condition depicted in Fig. 4D the body is orthogonal to Earth gravity and optimally oriented to detect wingbeat-driven modulation of G_z_. Head orientation, on the other hand, varies during each wingbeat cycle such that G_zH_ is not only attenuated compared to G_zB_ but its magnitude is asymmetric relative to G_zB_ at the acceleration peaks and troughs. That is, the attenuation in G_zH_ is greater at maximum downbeat acceleration compared to the attenuation at maximum acceleration at wing upbeat. This asymmetry in G_zH_ relative to G_zB_ is distinguished from that characterizing the coupled condition illustrated in Fig. 4A, in which maximum upbeat G_zH_ was approximately equivalent to maximum upbeat G_zB_ (acceleration troughs). In this uncoupled condition, maximum upbeat G_zH_ is consistently attenuated relative to G_zB_.

Fig. 4E represents another condition of head and body uncoupling in which inflight body orientation changes during each wingbeat cycle while head orientation is stable in the position to more optimally detect modulation in G_z_ due to wingbeat. In this configuration G_zB_ is asymmetrically attenuated relative to G_zH_ at both the maximum wing downbeat and upbeat. As discussed above, it is distinguished from the condition depicted in Fig. 4C by the consistent attenuation in maximum upbeat G_zB_, whereas in the coupled condition G_zH_ and G_zB_ at maximum wing upbeat are similar.

The data in Fig. 3 clearly demonstrate that inflight G_zH_ exhibits attenuation in maximum acceleration at the peaks and troughs associated with maxima during wing downbeat and upbeat, and that the attenuation is asymmetric where the G_zH_ and G_zB_ differences are greater at the peaks than the troughs. These findings were consistent with the uncoupled condition represented in Fig. 4D. Consequently, measurements of inflight head and body kinematics support the conclusion that *R. aegyptiacus* execute behaviors that minimize head acceleration (and, consequently, head displacement) and result in head stabilization during stable flight.

### Association of tongue click onset with head kinematics suggests relationship of acoustic probing with wingbeat cycle

The analyses of inflight head and body kinematics indicated that the *R. aegyptiacus* subjects exhibited head stabilization behaviors during the downbeat phase of the wingbeat cycle. One interpretation of this behavior is that it limits head movement during echolocation probing of the environment to stabilize inflight sensory gaze (Eitan et al., 2019). If this is true, it might be expected that tongue clicks were emitted during this period of the wingbeat cycle and head stabilization behavior. The direct measurements of head movements and sonar emissions required to establish this correlation have not been previously explored. Therefore, recordings from the head tag ultrasonic microphone were analyzed to first identify the tongue click emissions, and then determine the precise time of first emission of the click-doublet. We then determined G_zH_ and G_zB_ at these times, and evaluated their distributions for each subject. These analyses are illustrated in the flight epoch and acoustic recording shown in Fig. 5A, demonstrating identification of the tongue click onset (yellow diamonds superimposed on the acoustic recording, inset *Pressure (Pa)* vs *Time (s)*) and estimated G_zH_ (by interpolation; see *Methods*) at these times. Of the 50 flight epochs evaluated in both *R. aegyptiacus* subjects (25 each), a total of 525 click emissions were identified (i.e. *Red*: n=197; *Blue*: n=328). The distributions of G_zH_ at click onset for *Red* and *Blue* subjects are shown in Fig. 5B as box-and-whisker plots, illustrating that G_zH_ was ≥1g at the time of the first click for approximately 75% of the clicks in both subjects. The median G_zH_ values for the distributions from *Red* and *Blue* were 1.39g and 1.43g, respectively. As shown in Figs. 2 and 5, G_zH_ measures ≥1g correspond to wing downbeat. In view of the similarity in this behavior in both subjects, the data were combined to produce a single distribution of first-click G_zH_ measures (n=525), for which the median was found to be 1.41g.

**Figure 5.**
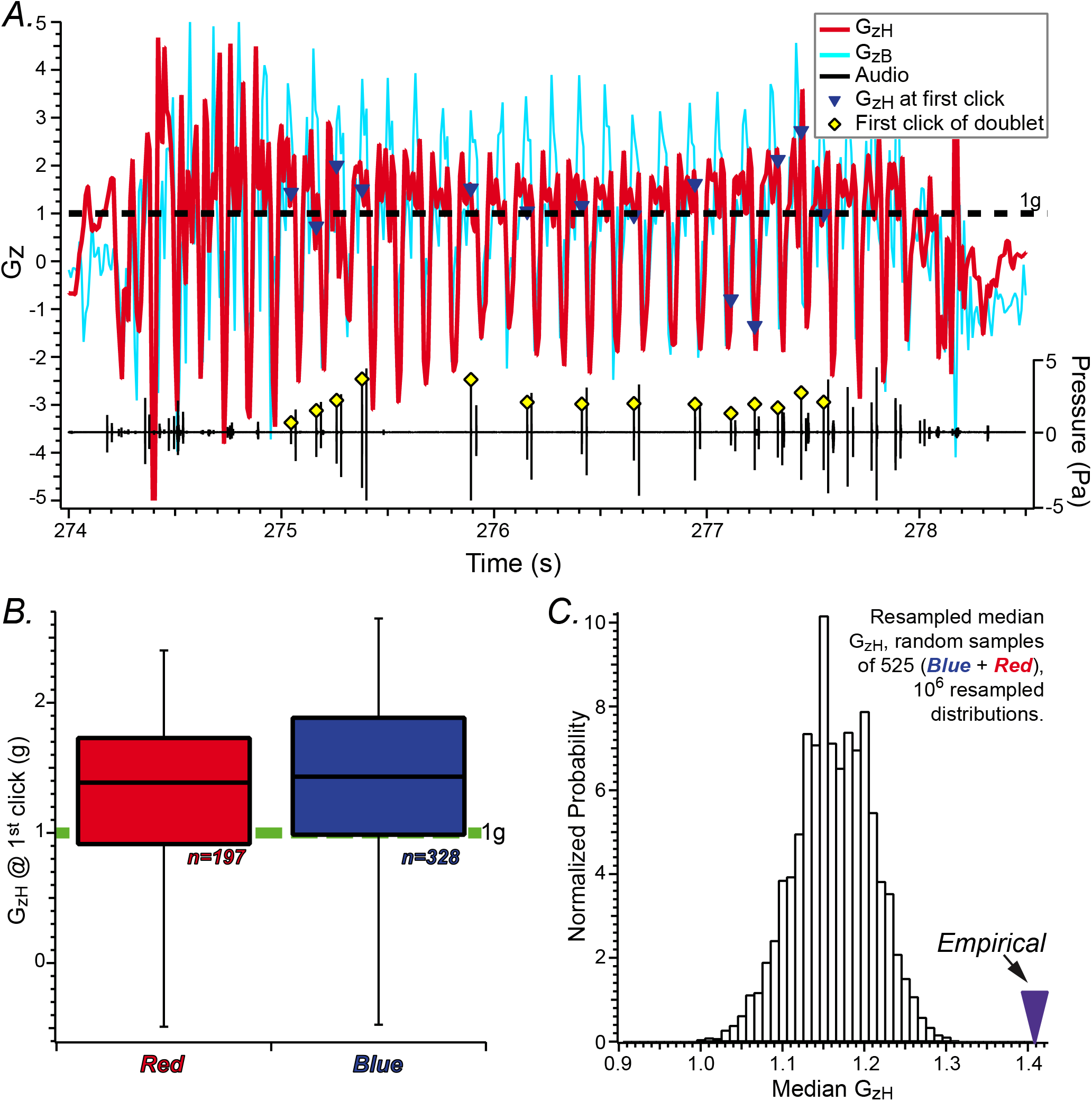
Onset times of tongue clicks exhibit greater likelihood of occurring at G_zH_ > 1 and the period of head stabilization. *A.* Representative record of GzH, G_zB_, and the associated audio recording exhibiting representative tongue click doublets. These data illustrate the identification of tongue click onsets shown by the red-filled diamonds superimposed on first click of the doublets within the selected period of stable flight. The corresponding values of G_zH_ at these times are identified by the inverted blue triangles. For the 14 identified onset times, 10 corresponded to G_zH_ values > 1g (71%). *B.* The distributions of G_zH_ at first click onsets for each of the *R. aegyptiacus* subjects are shown as box-and whisker plots, illustrating the similarity in their distributions. The data from both subjects were combined to test the null hypothesis that the median G_zH_ value from all G_zH_ measures corresponding to first click onset times could be derived from random association of onset time with GzH. *C.* The histogram represents median G_zH_ resulting from 10^6^ resampled distributions of 525 random association between click onset and GzH. The empirical median is shown as the inverted triangle at approx. 1.41g, indicating the value of the empirical median could not be derived from 10^6^ random distributions. This finding indicates that the probability of achieving the empirical median G_zH_ from randomly associated distributions is less than 10^−6^, supporting rejection of the null hypothesis.

To further evaluate whether the association between G_zH_ and first-click of a sonar doublet could be associated with head stabilization behavior, we tested the null hypothesis that the median G_zH_ from the combined 525 first-click events could have been derived from random association of click onset times and G_zH_. This was done by computing the median G_zH_ measure from a distribution of 525 random G_zH_ samples obtained from flight epochs in both *R. aegyptiacus* subjects. This was repeated one million times to produce 10^6^ median values, the distribution of which is shown in the histogram of Fig. 5C along with the median value from the empirical first-click G_zH_ measures. This analysis demonstrates that the probability of obtaining the empirical median G_zH_ (i.e. 1.41g) from random associations of G_zH_ and click onset was less than 10^−6^, supporting rejection of the null hypotheses. Note that the distribution of random G_zH_ medians ranged from 1.0 – 1.3, reflecting the bias of inflight G_zH_ >1g. A comparable resampling analysis was also determined for G_zB_ (not shown), producing a similar result as for G_zH_ wherein the probability of deriving the empirical G_zB_ median from a random association of G_zB_ and click onset was less than 10^−6^ and supporting rejection of the null hypothesis. If the associations between G_zH_ and G_zB_ and click onset were not random, they must be due to underlying factors driving the associations.

## Discussion

### Echolocation probing during wing downbeat

Previous investigations have interpreted head stabilization behaviors during locomotion as mechanisms to optimize sensory gaze (Brooks and Cullen, 2019; Dietrich et al., 2020; Dietrich and Wuehr, 2019b; Eitan et al., 2019; Goldberg and Cullen, 2011; Kress et al., 2015). If similar mechanisms apply to acoustic gaze stabilization during echolocation, it would be expected that inflight emission of sonar signals would be temporally associated with periods of head stabilization. The data for inflight head and body kinematics revealed evidence for head stabilization in *R. aegyptiacus* during periods where G_zH_ > 1 (Fig. 3 and 4) and wing downbeat. Therefore, the onset of sonar doublets emitted by *R. aegyptiacus* during periods where G_zH_ > 1 would strongly infer the implementation of head stabilization mechanisms to enhance acoustic gaze. Through resampling analyses we tested the (null) hypothesis that the empirical distribution of G_zH_ at click onset could be derived from random association of first click times and G_zH_ (Fig. 5C). We found this probability to be extremely low (p < 10^−6^), supporting rejection of the null hypothesis and supporting the existence of an underlying mechanism responsible for the temporal association of head stabilization and echolocation emissions. Therefore, these analyses are consistent with the notion that *R. aegyptiacus* execute head movement behaviors to stabilize acoustic gaze during exploratory flight.

The correlation between onset of echolocation emission and wingbeat has been established in many laryngeal echolocating species (Holderied and von Helversen, 2003; Jones, 1994; Kalko, 1994; Suthers et al., 1972; Wong and Waters, 2001; Yartsev and Ulanovsky, 2013). In laryngeal echolocators emissions were coupled to expiration and wing upbeat, and it was proposed that this may be driven by energy efficiencies (Holderied and von Helversen, 2003; Jones, 1994; Kalko, 1994; Suthers et al., 1972; Wong and Waters, 2001). However, echolocation appears to have minimal energetic costs (Speakman and Racey, 1991), even under various flight conditions with higher energy demands (Voigt and Lewanzik, 2012). A more recent analysis provided evidence that high intensity echolocation calls did, in fact, impose high metabolic demands (Currie et al., 2020), suggesting that energy efficiencies under these conditions may be derived by coupling sonar calls to expiration and wing upbeat. Yartsev and Ulanovsky (2013) previously showed that echolocation signal production was correlated with wingbeat in *R. aegyptiacus*, a lingual echolocator. However, because lingual echolocation is not driven by expiratory activity, it is unlikely that the coupling of echolocation signal production and wingbeat would offer any energetic advantage as may occur in laryngeal echolocators. This rationale further supports the conclusion that the temporal correlation of sonar onset and wingbeat optimizes acoustic gaze stabilization in *R. aegyptiacus*. If acoustic gaze stabilization is a sensorimotor imperative for all echolocating bats, it may be expected that sonar call emission would also be associated with head stabilization behaviors in laryngeal echolocating species. This suggests that, in these species, head stabilization behaviors would be associated with wing upbeat, an important hypothesis that is imminently testable.

### Head stabilization may be dependent upon behavioral state

The direct measurements of G_zH_ during exploratory flight from *R. aegyptiacus* obtained in the present investigation demonstrate that it exhibited wingbeat-cycle modulation of at least 3g (estimated from the difference in median peak and trough GzH, Figs. 3C and D). At wingbeat frequencies of approximately 8Hz this G_zH_ range corresponds to peak-to-peak displacements of approximately 8mm. In their experiments tracking head and body movements during an approach task to a landing target, Eitan et al. (2019) found that *R. aegyptiacus* exhibited dramatically attenuated inflight vertical gaze angles compared to body-target angles. This was interpreted to represent head stabilization behaviors serving acoustic gaze optimization during this task. The data on vertical gaze angle (determined from continuous tracking of head position relative to the fixed target) did not appear to exhibit similar wingbeat-cycle modulation as body-target angle, and even exhibited periods of very little deviation. Despite the evidence of head stabilization behaviors demonstrated by direct recording of head and body kinematics, the periods of stable flight examined in the present analyses exhibited robust modulation of head acceleration phase-locked to body acceleration and wingbeat cycle (Figs. 3 and 5). Therefore, it appears that head stabilization behaviors during the target-approach task tested by Eitan and colleagues (2019) may be more refined than those exhibited during periods of stable flight investigated in the present study. A direct comparison of the head displacement magnitudes under the two flight conditions was not readily achievable in view of the different measurement parameters offered by Eitan et al. (2019) and the present study. Such a comparison would be invaluable to better understand heterogeneities in behavioral state-dependent acoustic gaze stabilization.

### Could inflight kinematics be influenced by loads imposed by head- and body-mounted tags?

The use of miniature body-affixed sensors to measure various attributes of natural behaviors broadly expands the repertoire of documentable characteristics by enabling high resolution measurement capabilities unrestricted by either cabling or video documentation. The potential disadvantage is that they impose additional loads to the experimental subjects, which could potentially affect the behaviors under measurement. As previously noted, Stidsholt et al. (2018) found no evidence that adding a single 2.6-gram motion tag influenced the flight characteristics of two bat species for which the tags added as much as 20% of the animals’ body mass. Similarly, if the flight behaviors of the *R. aegyptiacus* subjects of the present study were influenced by the imposed 5.2 grams of additional load (combined mass of head and body tags) it might be expected that the flight characteristics of *Blue* and *Red* (163 and 218 grams body mass, respectively) would differ in view of the 50 gram difference in body mass. However, the flight characteristics exhibited by both subjects during exploratory flight were generally similar (Fig. 3), and the median wingbeat frequencies were virtually identical. These observations support the contention that affixing the motion tags to the head and body had no noticeable impact on their flight behaviors.

Other investigations employing body-mounted tags on larger bats (*Rhinopoma microphyllum*, 40 – 45 grams, carried tags amounting to >5% body mass (Cvikel et al., 2015); *R. aegyptiacus*, approx. 180 grams, carried tags amounting to approx. 12% body mass (Danilovich et al., 2015)) arrived at similar conclusions. Egert-Berg et al. (2018) tagged 5 species with devices (microphone and GPS) representing up to 15% additional loads as well as lighter tags, and concluded that additional loading represented no deleterious effects on flight and foraging behavior. These data suggest that inflight behaviors of most bats are very robust to additional loads.

While several investigations utilized body-mounted instrumentation, there is a paucity of information regarding the impact of head-mounted hardware on head and body kinematics in flying bats. Interestingly, Yartsev and Ulanovsky (2013) found that the dominant wingbeat frequency in *R. aegyptiacus* subjects carrying a 12 grams head load was approximately 8Hz, similar to the wingbeat frequencies exhibited by subjects of the present study. Despite differences in head and body loading represented in the two investigations, these data suggest that general flight characteristics of *R. aegyptiacus* are not dramatically impacted by the passive loads imposed in these experiments.

Despite the stability in general flight characteristics to modest passive loads, the impact of the 2.6 grams head load imposed by the motion tag upon measures of head kinematics was further scrutinized. Unfortunately, the present dataset was not sufficient to unequivocally exclude the possibility that the attenuation in peak G_zH_ during wing downbeat was influenced by passive load imposed by the motion tag. However, multiple lines of evidence suggested that any effect of passive head loading would be modest. Non-instrumented *R. aegyptiacus* subjects exhibited robust head stabilization behaviors during target identification and landing tasks in the dark and without any head load (Eitan et al., 2019). The present study provides evidence that such behaviors observed under unloaded conditions extend to exploratory flight, indicating that head stabilization is an integral component of natural acoustic gaze stabilization and, therefore, not dependent upon head loads imposed in the laboratory. Furthermore, estimates of head mass in *R. aegyptiacus* indicated that it represented approximately 12% of total body mass, suggesting that head masses of the two subjects of the present study were 19 and 26 grams (*Blue* and *Red*, respectively). The 2.6 grams motion tags, then, imposed relative loads of 13 and 10% to the heads of these subjects. A wholly passive model of head stabilization in whooper swans, with necks of approximately equal length as their bodies, predicted a 16% decrease in head displacement *gain* (i.e. Δhead/Δbody) with a 20% increase in head load (Pete et al., 2015). This analysis generally predicts that, even under passive head stabilization conditions, the loads imposed by the motion tags in the present study would render a relatively small effect on head kinematics (i.e. < 10%).

Head loading imposed by the motion tags may not be the only anomalous factor with the potential to influence flight behavior in bats. An analysis of swimming behavior in dolphins showed that placement of a back-mounted tag caused the instrumented subjects to swim at slower speeds, and fluid dynamics modeling indicated this was a direct outcome of increased drag imposed by the tag (van der Hoop et al., 2014). To our knowledge comparable effects of increased drag on inflight bat behavior have not been analyzed. However, to the extent that wingbeat frequency is a critical prognosticator of flight behavior, and observations of similar wingbeat frequencies under the drags imposed by low- and high-profile tags used presently and by Yartsev and Ulanovsky (2013) respectively, it appears that any effects of tag-imposed drag were small. With increased applications of head- and body-mounted instrumentation to the study of flight behavior, however, a critical look into the potential effects of imposed drag and weight on flight behavior is warranted.

### Vestibular contribution to acoustic gaze stabilization

The neural mechanisms supporting sensory gaze stabilization during locomotion are not completely understood. Previous investigations provided compelling evidence that efference copy (i.e. “copies” of spinal locomotor pattern generator activity projecting rostrally to CNS circuits serving sensorimotor integration) plays an important role in visual gaze stabilization in late stage larval *Xenopus* (Chagnaud et al., 2015; Lambert et al., 2012). These circuits not only drive compensatory oculomotor behavior in the principal plane of locomotor-associated head movements (Lambert et al., 2012), but also were found to suppress sensory input representing locomotion-associated head movements from the peripheral vestibular epithelia (Chagnaud et al., 2015). The precise mapping of active movement suppression of peripheral vestibular input, however, may not generalize phylogenetically, as comparable suppression of afferent vestibular signaling during active movement was not found in primates (Brooks and Cullen, 2014). Dietrich and Wuehr (Dietrich and Wuehr, 2019a) recently reported data to potentially reconcile these differences, finding that during human ambulatory activity horizontal gaze remained dependent upon vestibular input while vertical gaze stabilization relied more on efference copy. The latter investigation supports the idea that gaze stabilization is “spatially tuned” to principal locomotor-associated head movements requiring stabilization.

Aerial locomotion in *R. aegyptiacus* appears to be similar to terrestrial locomotion exhibited by bi- and quadrupeds in that both are associated with robust head movements in the Earth-vertical plane (i.e. G_z_) ((Dietrich and Wuehr, 2019a); L.F. Hoffman, unpublished data). Hence, it might be expected that a principal driver of head stabilization in *R. aegyptiacus* is efference copy of wingbeat spinal pattern generators. This would be consistent with the notion posited by Eitan et al. (2019) that enhanced stabilization behavior exhibited by *R. aegyptiacus* as target distance decreased argued against the notion that it reflected compensatory mechanisms driven by vestibular input. However, vestibular input was shown to be critical for inflight obstacle navigation in echolocating bats (Horowitz et al., 2004). Furthermore, preliminary studies of the peripheral vestibular epithelia in *R. aegyptiacus* revealed cellular adaptations consistent with enhanced high-frequency coding capabilities by semicircular canal cristae compared to terrestrial rodents (L.F. Hoffman, unpublished data). These findings suggest an important contribution of peripheral vestibular mechanisms to natural behaviors of bats. Consequently, further investigation of the challenges of sensory integration during high-performance locomotor activity exhibited by bats will ameliorate our general understanding of the neural mechanisms underlying sensorimotor integration and gaze stabilization.

## Acknowledgements

The authors also acknowledge Mark Johnson for development of the motion tags that made this investigation possible.

